# Differential ability of three bee species to move genes via pollen

**DOI:** 10.1101/2022.07.08.499290

**Authors:** Fabiana P. Fragoso, Johanne Brunet

## Abstract

Since the release of genetically engineered (GE) crops, there has been increased concern about the introduction of GE genes into non-GE fields of a crop and their spread to feral or wild cross-compatible relatives. More recently, attention has been given to the differential impact of distinct pollinators on gene flow, with the goal of developing isolation distances associated with specific managed pollinators. To examine the differential impact of bee species on gene movement, we quantified the relationship between the probability of getting a GE seed in a pod, and the order in which a flower was visited, or the cumulative distance travelled by a bee in a foraging bout. We refer to these relationships as ‘seed curves’ and compare these seeds curves among three bee species. The experiments used *Medicago sativa* L. plants carrying three copies of the glyphosate resistance (GR) allele as pollen donors, such that each pollen grain carried the GR allele, and conventional plants as pollen recipients. Different foraging metrics, including the number of GR seeds produced over a foraging bout, were also quantified and contrasted among bee species. Leafcutting bees produced the lowest number of GR genes in a foraging bout, and moved them the shortest distances, bumble bees the longest. Values for honey bees were intermediate. Seed curves correlated with field-based gene flow estimates. Thus, differential seed curves of bee species, reflecting within foraging bout patterns, translated into distinct abilities of bee species to move genes at a landscape level. Differences in seed curves reflected differences in foraging behavior among bee species, and helped explain their differential impact on gene flow and the spread of GE genes in insect-pollinated crops.

## Introduction

Gene flow is an important evolutionary force, and high gene flow tends to homogenise the genetic diversity of plant populations [1-2]. In flowering plants, gene flow occurs predominantly through the dispersal of pollen, and subsequent seed set from pollinated ovules [3-4]. Gene flow can also occur through the dispersal of seeds [5-6]. Insects move pollen between visited flowers, and most flowering plants, including many crops (a large number of fruit crops, most vegetables when one considers seed production, together with some forage and oil producing crops), are insect-pollinated. As the agents responsible for the transfer of pollen from flower to flower in the majority of angiosperms, pollinators impact the movement of genes via pollen [7-9] and affect plant reproductive success [10-11].

In agriculture, gene flow can occur among fields of a crop or between a crop and a cross-compatible feral or wild relative [12-13]. The seed industry has long been interested in limiting gene flow among fields of a crop in order to maintain the genetic purity of crop varieties. Moreover, since the release of genetically engineered (GE) crops, there has been increased concern about the introduction of GE genes into non-GE fields of a crop, and their spread to feral or wild cross-compatible relatives [14-17]. The movement of pollen from a GE to a non-GE variety of the same crop can contaminate seed lots, create adventitious presence (unwanted GE genes in seed lots), and offset coexistence of different markets with serious negative economic impacts [18-19]. In fact, the organic and export markets have a very low tolerance threshold for GE genes and their presence can disqualify a seed lot for the intended market. In addition, in cross-compatible feral or wild relatives of a crop, the GE gene transmitted from the GE crop via pollen or seeds could increase in frequency with potentially negative consequences [13,16, 20-21]. For example, a gene that increases the competitive ability of the plant could increase the weediness and invasiveness of the wild or feral species and consequently reduce the biodiversity of the community [13,16]. The rapid development of genome editing technologies is expected to create an increase in both the numbers and acreages planted to GE crops, with increasing concern for gene flow to feral and wild populations, and for adventitious presence that can disrupt the coexistence of different seed markets (organic, conventional, export) and negatively impact the seed purity of cultivars. It is imperative to increase our understanding of how pollinators impact gene flow from GE crops.

A standard and efficient method of limiting gene flow to ensure cultivar purity and limit adventitious presence is via isolation distance [22]. It is becoming more evident, however, that the optimal isolation distance may vary with pollinator species, because gene flow can vary with bee species [23,24]. In alfalfa, for example, genes have been recovered farther distances in fields pollinated with honey bees, relative to fields pollinated with leafcutting bees [23]. These studies were done in different areas, and these findings have led to an isolation distance of 274m for leafcutting bees, and 4.8 km for honey bees [24]. In contrast with pollen dispersal, where pollen carryover curves have been described for distinct pollinators [25-28], we have little information linking gene flow to pollinator behavior (but see [9]). A pollen carryover curve describes the decay in pollen deposition as a pollinator moves from flower to flower along a foraging path [29]. While gene flow is related to pollen dispersal, not all pollen grains will set seeds, and therefore the shape of the pollen carryover curve could differ significantly from the pattern of seed set as a pollinator moves from flower to flower and travels increasing cumulative distance. The only study, to our knowledge, to examine dispersal of genes instead of pollen used paternity shadows which describe the distribution of progeny sired by pollen from a single donor flower [7]. While this approach helped parameterize a gene flow model [7], it is more difficult to relate to pollen carryover curves, and to link to pollinator behavior.

To examine dispersal of genes by distinct bee species, we used *Medicago sativa* L. plants carrying three glyphosate resistance (GR) alleles as pollen donors, and conventional (non-GR) *M. sativa* plants as pollen recipients. We described the relationship between the probability of getting a GR seed in a pod, and the order a flower is visited in succession by a bee or the cumulative distance it travels over a foraging bout. We refer to this relationship as a “seed curve” and quantify the “seed curve” for three bee species that pollinate *M. sativa*, the European honey bee, *Apis mellifera L*., the alfalfa leafcutting bee, *Megachile rotundata F*., and the common eastern bumble bee, *Bombus impatiens* Cresson. In addition, to further understand the pattern of GR seeds set by these three bee species, we contrasted different foraging metrics among bee species, such as the number of flowers visited and tripped, and the number of GR seeds per foraging bout. Finally, we examined how the gene flow predictions based on the seed curves, compared to gene flow estimates previously obtained in seed-production fields for these bee species. Understanding how seed curves vary among pollinator species can increase our understanding of why distinct pollinators have different impact on gene flow, and help determine whether some pollinator characteristics are linked to greater gene flow. Such knowledge would help biotechnology regulators as they refine the rules for isolation distances between fields pollinated by specific pollinators. The rapid development of novel genome editing technologies and the eminent release of new GE varieties, has heightened the importance of understanding how distinct bee species differentially affect gene flow, coexistence, and the spread of GE genes to sexually compatible feral and wild populations.

## Materials and Methods

### Study system

*Medicago sativa* L. (Fabaceae), known as alfalfa or lucerne, is a tetraploid, self-compatible, perennial legume, widely cultivated worldwide. Flowers are organized into clusters or racemes and range in color from pale to dark purple [11]. Alfalfa relies mainly on bees for pollination and subsequent seed set, and a tripping mechanism enables pollen transfer. When a pollinator presses the two lower petals that form the keel, the enclosed stamens and pistil are forcefully released, and pollen is deposited on the pollinator’s body. At the same time, pollen already on the pollinator’s body gets deposited on the stigma. Flowers that are not tripped do not set or sire seeds.

Managed pollinators in alfalfa seed production in the United States include the alfalfa leafcutting bee, *M. rotundata* and the European honey bee, *A. mellifera*. The strict habitat requirement of the alkali bee, *Nomia melanderi* Cockerell, restricts its use to the Walla Walla valley in Washington [24]. Wild bees, including the common eastern bumble bee, *B. impatiens*, also visit alfalfa flowers [30]. Although bumble bees are used as pollinators in alfalfa breeding programs, they are not used in alfalfa seed production in the United States, where wild bumble bees do not play an important role. These distinct bee species trip flowers at different frequencies (tripping rates) [9, 30], and a higher tripping rate is linked with a greater seed set per bee visit [11]. While alfalfa leafcutting bees have high tripping rates and are efficient pollinators, honey bees learn to avoid the tripping mechanism and often become nectar robbers by drawing nectar from the side of the flower.

### Experimental Design

The experiments were carried out inside cages set up in a greenhouse room at the University of Wisconsin Walnut Street Greenhouse in Madison, WI, USA. A smaller 2.0 × 2.0 × 1.8 m cage adjoined a larger 4.0 × 2.0 × 1.8 m cage (Fig 1). The frames of the cages were made of PVC tubing and were covered with mosquito netting (Skeeta, Inc., Bradenton, FL, USA) which was also used to separate the two adjacent cages. Glyphosate-resistant (GR) alfalfa plants, also known as Roundup Ready (RR) alfalfa, were placed in the small cage and used as pollen donors in the experiment. These plants had at least three copies of the glyphosate-resistant allele to ensure that all pollen grains received at least one copy. Conventional alfalfa plants were placed in the large cage and were used as pollen recipients.

**Fig 1.**
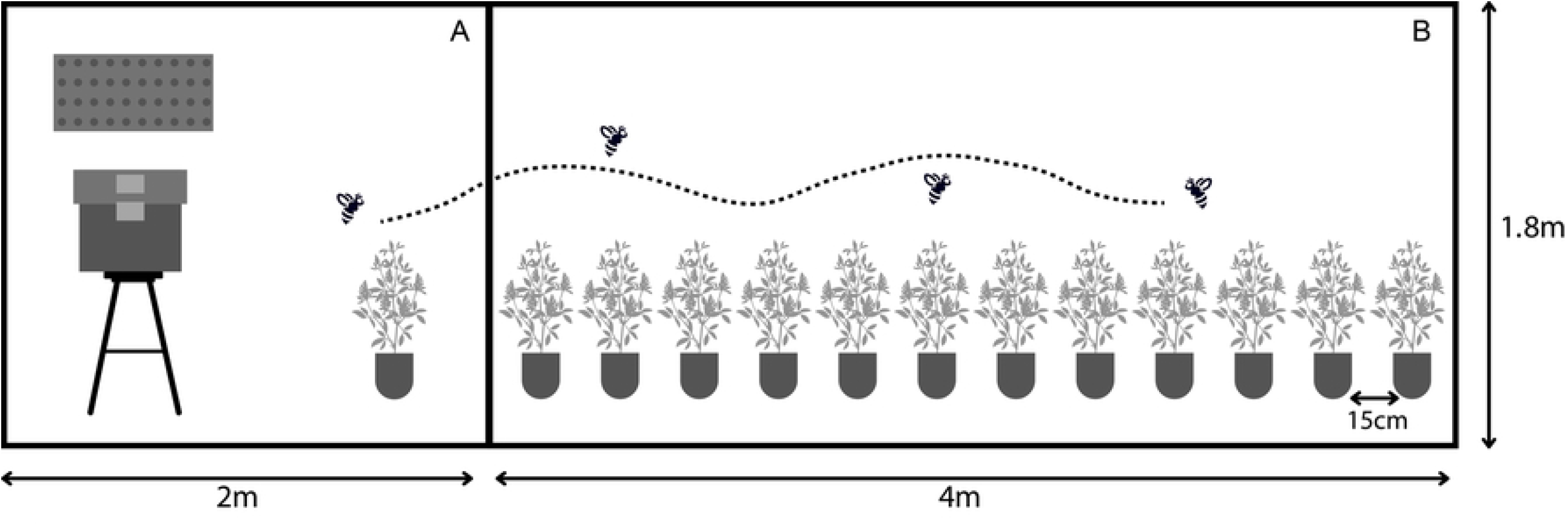
Experimental set up. Bee hive, or nesting board with bees, were kept in a small cage (A) adjacent to a large cage (B). Prior to trials, bees were trained in the small cage on conventional alfalfa plants. During a trial, glyphosate-resistant (GR) donor plants were first presented to a bee inside the small cage. After tripping five flowers on donor plants, the bee was moved to the large cage to forage on conventional plants.

Trials were run for one bee species at a time. Prior to starting a set of trials, bees were trained to visit alfalfa plants in the small cage, using several blooming conventional alfalfa plants. For bumble bees, we used a hive containing 75-100 workers (Koppert Biological Systems, Howell, MI, USA) and for honey bees, a four-frame nucleus honey bee colony (Gentle Breeze Honey, Inc., Mount Horeb, WI, USA). For leafcutting bees, 30-50 bees, incubated from commercially available cocoons, were released into the training cage every two days for the duration of the experiment. Both male and female leafcutting bees were released in the small cage to allow mating, because mated females collect pollen and nectar to provision their eggs. Only female bees were used in the trials. A 30 × 50 cm nesting board was placed in the small cage to permit leafcutting bees to nest. The hive or nesting board was set on the side of the small cage opposite the opening to the large cage (Fig 1).

Trials began after bees foraged consistently on alfalfa in the training cage. Prior to starting a trial, training alfalfa plants were removed from the small cage and bees were not allowed to visit any plant for at least 15 hours. We could close the bumble bee hive the evening prior to a trial, but honey bees and leafcutting bees remained free inside the cage. However, in the absence of plants, most honey bees would stay inside the hive and leafcutting bees would lay still on the netting. To set up a trial, two flowering GR alfalfa plants, used as pollen donors, were placed in the small cage. In addition, 12 conventional flowering alfalfa plants, used as pollen recipients, were placed in a linear array in the larger cage, each plant separated by 15 cm (Fig 1). Floral display size on recipient plants was standardized to 5-6 racemes per plant and 4-6 untripped flowers per raceme. A thin thread of a different color was used to distinguish the five racemes on each recipient plant, and each flower within a raceme was marked with a small dot of a distinct colored paint. Based on our observations, the markings did not affect bee behavior.

### Bee observations

Only one individual bee was tested in the large cage at any given time. For bumble bees, one individual bee was released from the colony into the small cage. After the bumble bee tripped five GR flowers from the donor plants in the small cage, the screen separating the cages was lifted and the bee was moved to the larger cage. Honey bee and leafcutting bee remained free to forage inside the small cage, and the first honey bee or female leafcutting bee to start foraging on the donor plants in the cage was tracked until it tripped five flowers. The honey bee or female leafcutting bee was then caught in a centrifuge tube and moved to the large cage. Tripping five flowers required visiting, on average, between six and 14 flowers, depending on the bee species.

As soon as a bee started foraging on the conventional plants in the large cage, we used a voice recorder to note each plant, raceme and flower the bee visited in succession, from the beginning to the end of the foraging bout. A foraging bout could start from any plant position and revisits to plants and flowers were permitted. For the social bees, a foraging bout ended when the bee started flying back towards the hive, and for the leafcutting bees a bout ended when the bee landed for more than two minutes without grooming. Prior experience indicated that leafcutting bees were unlikely to keep foraging after this two-minute rest period.

### Flower order, distance travelled between successive flowers, and fruit collection

Each flower visited in succession during a foraging bout was given a flower order corresponding to the order in which it was visited. At the end of each trial, we measured the distance between all pairs of flowers visited in succession by a bee. Each tripped flower was marked with a numbered tag and periodically checked for pod (fruit) development. Untripped flowers do not form pods because they remain closed, with no pollen being removed from the anthers or deposited on the stigmas. Pods were collected 4-6 weeks following a trial, when seeds were mature. Each individual pod with the associated tag was kept in a separate coin paper envelope and transferred to a refrigerator until pods could be hand-threshed to extract the seeds, and seeds could be counted and tested for the presence of the glyphosate resistance gene.

### Detection of the glyphosate resistance gene

Seeds collected from the bumble bee and leafcutting bee trials were tested for the presence of the glyphosate resistance gene using a seedling phenotypic assay, combined with test strips [31]. The phenotypic assay is a modification of a protocol originally developed by [32] to distinguish between conventional and glyphosate-resistant seedlings. The test strips, lateral flow immunoassay AgraStrip® RUR Seed and Leaf TraitChek™ (Romer Labs Inc., Union, MO, USA), detect the CP4 EPSP protein. Because the incubator needed to grow the seedlings was not accessible during the COVID-19 stay at home period, seeds collected from the honey bee trials were all individually tested using the test strips.

For the phenotypic assay, seeds were placed in petri dishes lined with filter paper and watered with a low-concentration (80 ppm) glyphosate solution. The petri dishes were placed in an incubator set at 20°C, under a 16h light: 8h dark regime for 14 days. Three days following the initial set-up, seedlings were watered again with the low-concentration glyphosate solution. Thereafter, seedlings were watered with a 0.1% Plant Preservative Mixture (PPM, Caisson Laboratories, Inc., Smithfield, UT) deionized water solution to prevent fungal growth. At the end of the 14-day period, seedlings were evaluated for traits associated with glyphosate resistance, such as increased root length, smaller root width, and development of secondary roots or root hair [31-32]. The presence of the glyphosate resistance gene in each seedling categorized as positive using the phenotypic assay was confirmed using a test strip, following [17]. Seedlings scored as conventional based on the phenotypic assay were bulked in groups of twenty before applying the test strip. When a positive result was detected in a bulked sample, each seedling was individually tested with a test strip using its remaining cotyledons.

### Model selection

We used a binomial error distribution because our data structure included binomially distributed constrained counts (presence or absence of a GR seed). We examined the logit of the probability of getting GR seeds in a pod (log [p/(1-p)], with increasing order of flower visited or cumulative distance travelled in a foraging bout, with generalized linear mixed models (GLMM). We used a logit transformation for the dependent variable, log [p/(1-p)], to limit the values of p between 0 and 1. Here, p is the probability of getting a GR seed in a pod or the modelled proportion of GR seeds in a pod and (1-p) is the probability of getting a conventional (non-GR) seed in a pod. The origin for distance was the first flower visited and we added incrementally the distances between pairs of flowers visited in succession during a foraging bout. The model was fitted to the combined runs for each bee species using the “glmer” function in the lme4 package [33] in R [34]. The first model fitted to the data assumed a linear relationship between the logit of the dependent variable and each predictor (independent) variable (order of flower visited in a foraging bout or cumulative distance travelled). Because the logit of the probability of getting GR seeds in a pod may exhibit a different rate of decline as a function of the independent variable, we evaluated two additional models. The second model used the natural log of flower order or distance travelled, and the third model utilized the square root of flower order or distance travelled. In these two models, the relationship between the logit of the dependent variable and the independent variable is no longer linear. These models are abbreviated in the remainder of the manuscript, as the (1) “standard”, (2) “log”, and (3) “sqrt” models.

For each bee species and each predictor variable (flower order or cumulative distance), we fitted six different models. These models comprised the three types of models just described, standard, log or sqrt, to which we added either intercept or both intercept and slope as random effects. Each model has a fixed (population means) intercept and slope. In addition, the random intercept corresponds to a bee random effect and represents variation among individual bees in the logit of the probability of getting GR seeds set in the first flower visited in the foraging bout. The random slope considers variation among bees in the linear decline in the logit of the probability of getting GR seeds in a pod with increasing flowers visited or distances travelled. The six models per predictor variable per bee species were compared using the Akaike Information Criterion (AIC). The model with the lowest AIC was considered the best fit to the data and models were considered distinct if the difference in AIC between models was greater than two [35]. We examined a total of 36 models, two predictor variables times three bee species times the six models described above. The figures, presented for the best models, illustrate the back transformed logit, i.e. p or the probability of getting GR seeds in a pod. Logits were back transformed using the following equation: p = exp (a + b predictor variable) / [1 + exp (a + b predictor variable)]. Individual data points on the figures represent the observed proportions of GR seeds in a pod.

Analyses were performed on tripped flowers because untripped flowers do not set seeds. The order of visit of tripped flowers was determined and cumulative distance calculated between tripped flowers visited in succession. The cumulative distance was converted from centimetres to metres to facilitate model fitting. Because a flower is only tripped once, if a flower was visited more than once, we assumed it got tripped during the first bee visit. A previous study indicated no increase in the number of pollen grains deposited over repeated visits [29].

### Comparing foraging metrics among bee species

We compared different foraging metrics among the three bee species using one-way analysis of variance (ANOVA), followed by multiple pairwise-comparisons with Tukey HSD tests in R [34]. These foraging metrics included the total number of flowers visited per foraging bout, the proportion of flowers visited in a foraging bout that were tripped, foraging bout duration, and the cumulative distance travelled per foraging bout. A foraging bout starts with the first flower a bee visits in the array, and ends when the bee leaves the array. We also determined the proportion of flowers tripped in a foraging bout that set seeds, and the total number of seeds set and of GR seeds set per pod, and per foraging bout.

## Results

### Bumble bee models

We had 12 foraging bouts (runs) and 322 observations (pods with seeds). The logistic regression analysis only considers the pods that set seeds. For both flower order and cumulative distance, the log-odds ratio or logit of getting GR seeds in a pod, log_e_ (p/(1-p), was best described by the log model with random intercept (Table 1). For flower order, log_e_ (p/(1-p)) = 0.80 - 0.65 log_e_ flower order, with standard error = 0.53 for the intercept and 0.09 for the slope (Table 2). The standard errors (SE) around the slope or intercept for the fixed effects of the model (population means) illustrate the accuracy of the parameters. The standard deviation (SD) associated with the random effect for the intercept was 1.41 and illustrates variation among bees in the logit of getting GR seeds in the first flower visited in the foraging bout. For distance, the logit of getting GR seeds in a pod = -1.13 - 0.33 log_e_ distance with SE = 0.47 for the intercept and 0.05 for the slope. The standard deviation for the random intercept was 1.50. For both flower order and distance, the probability of getting GR seeds in a pod decreased with successive flowers visited or increased cumulative distance travelled by a bee (Figs 2 A and B).

**Table 1.** Comparing flower and distance models for three bee species.

| Bee species | bumble bee | honey bee | leafcutting bee |
| --- | --- | --- | --- |
|  | (N = 322) | (N =268) | (N =202) |
| **Flower Models** |  |  |  |
| Standard random intercept | 853.2 | **581.2** | 519.7 |
| Standard random slope and intercept | 840.8 | 584.6 | 511.5 |
| Log random intercept | **818.0** | 590.7 | 524.1 |
| Log random slope and intercept | 821.8 | 594.2 | 513.9 |
| Square root random intercept | 833.8 | **579.6** | 517.7 |
| Square root random slope and intercept | 828.9 | 583.1 | **508.1** |
| **Distance Models** |  |  |  |
| Standard random intercept | 861.1 | 585.8 | 523.3 |
| Standard random slope and intercept | 840.2 | 587.0 | 504.7 |
| Log random intercept | **821.3** | 613.7 | 542.2 |
| Log random slope and intercept | 823.8 | 616.5 | 537.3 |
| Square root random intercept | 839.9 | **581.8** | 513.4 |
| Square root random slope and intercept | 826.0 | 585.8 | **498.7** |

The models describe the logit of getting glyphosate-resistant (GR) seeds in a pod with increasing order of flower visited or cumulative distance travelled in a foraging bout for three bee species. A foraging bout starts with the first flower a bee visits and ends when the bee leaves the array. The
models with the lowest AICs are in bold. The sample size N represents the number of observations or the number of tripped flowers with seeds over all foraging bouts for each bee species.

**Table 2.** Fixed effect estimates for the best model(s) for flowers and distances for bumble bees (BB), honey bees (HB), or leafcutting bees (LCB).

| Bee Species | Best Model | intcp | SE | slope | SE |
| --- | --- | --- | --- | --- | --- |
| **Flower models** |  |  |  |  |  |
| BB | Log RI | 0.80 | 0.53 | -0.65 | 0.09 |
| HB | Standard RI | -0.45 | 0.25 | -0.04 | 0.00 |
| HB | Square root RI | 0.22 | 0.28 | -0.34 | 0.04 |
| LCB | Square root RIRS | 1.45 | 0.56 | -0.71 | 0.23 |
| **Distance models** |  |  |  |  |  |
| BB | Log RI | -1.13 | 0.47 | -0.33 | 0.05 |
| HB | Square root RI | -0.08 | 0.28 | -1.01 | 0.12 |
| LCB | Square root RIRS | 0.75 | 0.38 | -2.16 | 0.54 |

The models describe the log odds ratio (logit) of getting glyphosate-resistant seeds in a pod with increasing order of flower visited or cumulative distance travelled. The model parameters include the intercept (intcp) and the slope, with associated standard errors (SE). The variable RI stands for random intercept and RIRS for random intercept and random slope.

**Fig. 2.**
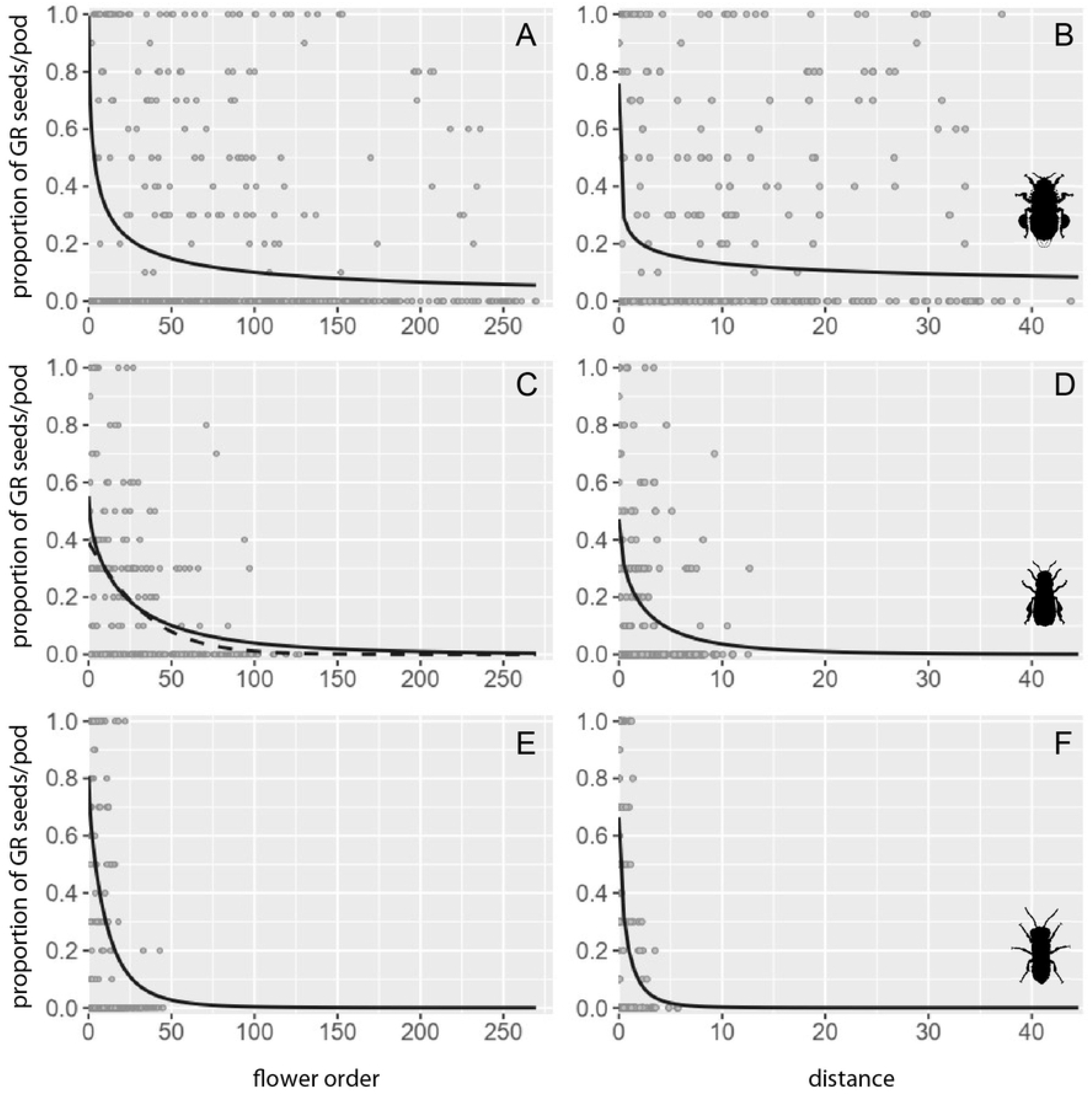
Probability of getting GR seeds in a pod with increasing flower order (left) or distance travelled (meter) (right) by a bee. Probabilities represent back transformed logits (see text for details). For bumble bee (upper plots), the log model with random intercept was the best model for both (A) flower order and (B) cumulative distance; for honey bee (middle plots), the square root (solid line) and the standard (dashed line) models with random intercept were equally good candidates for (C) flower order while the square root model with random intercept was the best model for (D) distance; for leafcutting bee (bottom plots), the square root model with random slope and random intercept was the best model for both (E) flower order and (F) distance.

### Honey bee models

The data for honey bees included 14 foraging bouts and 268 observations. We obtained two best models for flower order. The square root model with random intercept, and the standard model with random intercept (ΔAIC < 2) (Table 1). The equation for the square root model was log_e_ (p/(1-p)) = 0.22 - 0.34 sqrt flower order, with SE = 0.28 for the intercept, and 0.039 for the slope (Table 2). The standard deviation associated with the random intercept was 0.75. The slope and intercept for the standard model are presented in Table 2. For distance, the logit of finding GR seeds in a pod was log_e_ (p/(1-p)) = -0.08 -1.01sqrt distance with SE = 0.28 for the intercept and 0.12 for the slope (Table 2). The standard deviation associated with the random intercept was 0.82. In all cases, the probability of getting GR seeds in a pod decreased with increasing flower order or cumulative distance travelled. (Fig 2C and D).

### Leafcutting bee models

We obtained 27 foraging bouts and 202 observations for leafcutting bees. The square root model with random slope and random intercept best explained the logit of getting GR seeds in a pod, for both successive flowers and for cumulative distance (Table 1). For flower order, the equation was log_e_ p/(1-p) = 1.45 - 0.71 sqrt flower order, with SE = 0.56 for the intercept and 0.23 for the slope (Table 2). The standard deviations were 2.12 for the random intercept, and 0.89 for the random slope. For distance, the equation was log_e_ (p/(1-p)) = 0.75 - 2.16 sqrt distance, with SE = 0.38 for the intercept and 0.54 for the slope (Table 2). The standard deviation associated with the random intercept was 1.51 and with the random slope 1.90. The probability of getting GR seeds in a pod decreased with successive flowers and with cumulative distance travelled (Figs 2 E and F). The logistic models for leafcutting bees were the only models where the random slope variable, i.e. considering among bee variation in slope, improved the fit of the model to the data.

### Comparing models among bee species

Because the type of the best model (standard, log or sqrt), for either flower order or distance, differed among bee species (Table 2), we could not statistically compare the intercepts and slopes among bee species. However, to address potential differences among bee species, we can visually contrast the graphs illustrating the probability of finding a GR seed in a pod with successive flowers visited and cumulative distance travelled (Fig 2). Bumble bees had a fairly steep rate of decay in the probability of getting a GR seed in a pod (slope), but the probability levelled off after visiting approximately 30 flowers or traveling five meters, creating a distribution with a long tail (Fig 2A-B). When bumble bees moved pollen from flower to flower, GR seeds could still be found with a low probability in a pod, after a bee visited 250 flowers or travelled over 40 meters. Honey bees were intermediate between bumble bees and leafcutting bees, with a negligible probability of getting a GR seed in a pod after a honey bee visited approximately 100 flowers or travelled around 15 meters (Fig 2C-D). The rate of change in the probability of getting a GR seed in a pod with increasing flower order or distance was steep when pollen was moved by leafcutting bees (Fig 2E-F). The probability of getting a GR seed in a pod was negligible after a leafcutting bee visited 50 + flowers or travelled five + meters (Fig 2E-F).

### Foraging metrics

All foraging metrics, except the proportion of tripped flowers that set seeds, varied among bee species (Table 3). Bumblebees visited more flowers, spent more time foraging, and travelled greater cumulative distance in a foraging bout relative to honey bees or leafcutting bees (Table 3). Leafcutting bees tripped the most flowers and set a greater proportion of GR seeds per foraging bout relative to honey bees and bumble bees (Table 3). Honey bees set the most mature seeds per pod and were otherwise similar to either bumble bees or leafcutting bees, depending on the variable examined (Table 3). Although leafcutting bees set a greater proportion of GR seeds per foraging bout, they set fewer seeds per foraging bout, such that overall leafcutting bees set the lowest number of GR seeds during a foraging bout (Mean ± SE) (9.22 ± 1.31), relative to bumble bees (31.08 ± 11.87), and honey bees (17.07 ± 3.49). However, per pod, leafcutting bees set more GR seeds (1.47 ± 0.14) relative to honey bees (1.03 ± 0.20) and bumble bees (0.98 ± 0.31).

**Table 3.**
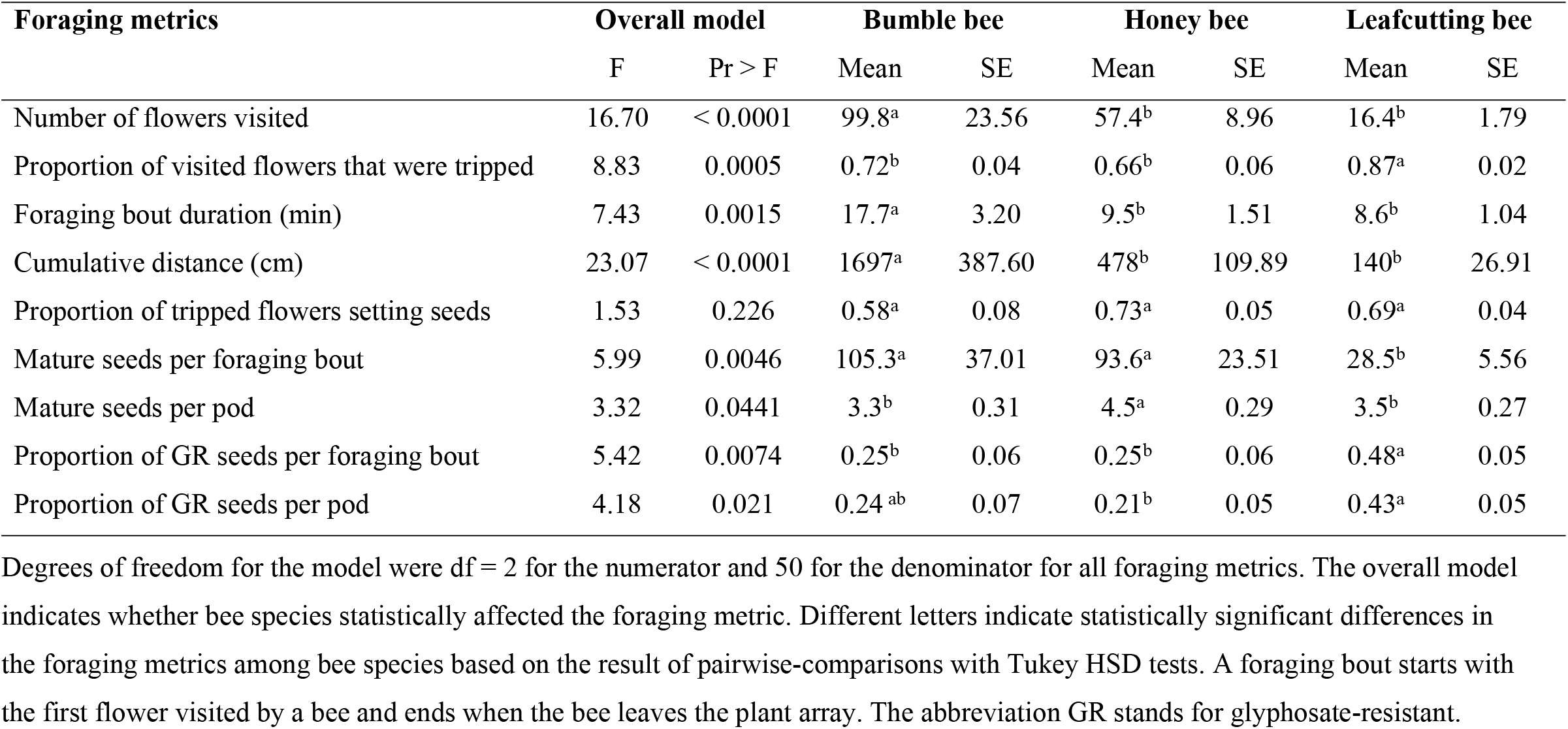
Differences in foraging metrics per foraging bout among bee species.

## Discussion

Leafcutting bees have the steepest rate of decay in the probability of setting a glyphosate-resistant seed, generate the lowest number of GR seeds per foraging bout (13.7), and carry these GR genes the shortest distances. Honey bees and bumble bees set similar number of seeds, and similar number of GR seeds per foraging bout (bumble bees: 26.3 GR seeds; honey bees: 23.4). Bumble bees have the dispersal curve with the longest tail, and moved more GR genes the longest distances. Honey bees moved genes intermediate distances. If we assign a probability of moving genes to each bee species, leafcutting bees would have the lowest probability, and bumble bees would have the highest.

While the current study examined the decrease in the probability of getting a GR seed in a pod over an average foraging bout, the findings are consistent with pollinator-mediated gene flow estimates obtained in alfalfa seed-production fields [23,36-38]. The gene flow estimates obtained from seed-production fields indicated lower distances travelled by GR genes when alfalfa leafcutting bees, relative to honey bees, were used as managed pollinators. In the USA, bumble bees are not used as managed pollinators in alfalfa seed-production fields, are not important pollinators in these fields, and we are not aware of gene flow estimates for bumble bees in seed production fields. For leafcutting bees and honey bees, predictions based on seed curves are consistent with gene flow estimates obtained in seed production fields [23]. This association is important because it supports a relationship between bee behavior within foraging bouts, and gene flow estimates at the field level. Such a connection is reminiscent of the relationships previously observed between behavior of birds observed on a small scale and seed dispersal at the landscape level [39-40]. These latter studies simulated the small-scale movements of birds, based on perching time, move length, and move direction, to predict bird-mediated seed dispersal of a plant species. Moreover, these results suggest how differences in seed curves among bee species can inform on the gene flow potential of these bee species. The observed differences in gene flow among bee species are further supported by the results of a 1965 experimental study that compared gene flow distances, using a white flowered alfalfa morph as pollen recipient, and variegated variety as pollen donor [41]. In this latter study, some bumble bee species were used and found to set seeds the farthest distances, followed by honey bees, and least leafcutting bees. Thus, gene flow predictions based on seed curves fit the observed gene flow data, linking gene flow estimates to bee behavior (seed curves). Next, we examine whether physical and behavioral differences among bee species help explain observed differences in seed curves.

Physical characteristics of bee species may affect gene flow. Previous studies have linked bee body sizes with their foraging ranges, defined as the maximum distance a bee species would travel from its nest or hive, based on homing, feeders training, or bee dance interpretation [42-44]. Larger bees fly longer distances although honey bees with their waggle dance have been predicted to forage farther distances relative to bumble bees despite their smaller size [45-47]. However, bee foraging range is quite different from the distance travelled while a bee is foraging for resources, and the latter measure is most likely to affect how pollen and the resulting seeds are spread [48]. In the current study, bumble bees, the largest of the three bees, visited the most flowers per foraging bout, and travelled the longest total cumulative distance. The smallest bees, the leafcutting bees, visited the least flowers, and travelled the shortest total cumulative distance. Honey bees were intermediate for both flowers visited and total cumulative distance travelled. The ranking of their body sizes correlates with the ranking of the distances travelled while foraging. In addition, a field study recorded bumble bees traveling the longest net distances as they foraged in alfalfa patches, followed by honey bees, while leafcutting bees travelled the shortest distances [9]. A net distance represents the distance between where a bee started and ended a foraging bout; it represents the straight-line distance between the first and last visited racemes. Future studies should increase the sample size of bee species to further strengthen the correlation. Available data indicate how bee body size is linked not only to foraging range, but also to the distance bees travelled while foraging, the latter measure being more relevant to pollen dispersal and gene flow.

Bee body size may also affect the size of the pollen load carried by the bee, which in turn can influence gene flow. Using radio frequency identification (RFID), Minahan and Brunet [49] determined that, despite spending similar amount of time during a foraging trip, bumble bees brought heavier pollen sacs back to the hive relative to honey bees. A foraging trip included the time between leaving and returning to the hive. Although pollen present in pollen baskets is not available for pollination, the heavier loads carried by bumble bees may also reflect a greater pollen load on the parts of their body that come into contact with plant stigmas. Bee species vary in the size of their pollen loads for specific plant species [50], and, in this respect, bumble bees deposited more GE pollen grains during an average foraging bout relative to leafcutting bees [29], and had more GE pollen grains left on their body following a foraging bout (unpublished data), suggesting greater pollen loads on their bodies. In addition, in the current study, bumble bees set the most GE seeds in a foraging bout, followed by honey bees, and least leafcutting bees, supporting a correlation between bee body size and the total number of GE seeds set in an average foraging bout. Bee body size is associated not only with bee movement [9], but also with pollen dispersal [29], and the resulting seeds (this study). It is also worth noting how a heavier pollen load on the bee’s body, combined with a larger number of flowers visited per foraging bout, and longer distances travelled, help explain the longer tail observed in the seed curve when bumble bees moved GE pollen, relative to the other two bee species.

The tripping rate, or the proportion of flowers visited by a pollinator whose anthers and stigmas were released from the previously closed flowers, has been proposed as a behavior affecting pollen dispersal and resulting seeds [9, 29]. Shorter pollen dispersal and distances of resulting seeds are predicted for pollinators with high tripping rates because pollen only gets deposited on tripped flowers [9]. Thus, the lower the tripping rate, the greater the number of untripped flowers visited, which increases the distance travelled by the pollinator before it deposits pollen on stigmas. For plant species without a tripping mechanism, a bee with a high tripping rate would be equivalent to an efficient pollinator. We thus expect efficient pollinators to move genes shorter distances. The tripping rate varies among the three bee species, with leafcutting bees typically tripping a greater proportion of visited alfalfa flowers relative to honey bees or bumble bees [9, 30]. Honey bees learn to avoid the tripping mechanism through experience, stealing nectar from the side, and naïve bees tend to trip more flowers than experienced bees [51]. However, the tripping rate of honey bees was quite high in the current study, likely because no other sources of pollen were made available and the bees had to gather their pollen from *M. sativa* flowers. In fact, in the current study the tripping rate of honey bees was similar to the tripping rate of bumble bees, which itself was higher than under field conditions (honey bee: 22.7 % and bumble bee: 50.7% over two years; [9]. While the high tripping rate of leafcutting bees could help explain its steep decline in the probability of setting a GR seed in a pod, tripping rate alone was not sufficient to explain the long dispersal tail of bumble bees.

Distinct pollinators have been previously shown to differentially affect plant reproductive success, pollen dispersal, and selection on floral traits [10-11, 29, 52-55]. Here, we show how distinct bee species have different seed curves, which predict relative gene flow by bee species observed in the field. In agriculture, a low probability of moving genes is associated with a low gene flow risk. Bees with lower gene flow risk such as leafcutting bees would generate the least adventitious presence, and minimize introgression and spread of GE genes to wild or feral cross-compatible populations. In natural populations, pollinators with a high probability of moving genes such as bumble bees would be linked to a greater impact on homogenising the genetic diversity of natural plant populations, and thus limiting genetic differentiation. Distinct bee species will have differential impact on the spread of GE genes in agriculture, and on the genetic differentiation of wild populations.

## Conclusions

Based on their seed curves and total number of GR seeds produced within foraging bouts, leafcutting bees have the lowest probability of moving genes, and moves genes the shortest distances; honey bees are intermediate; and bumble bees move genes the farthest and set the most GR seeds per foraging bout. Interestingly, the differential seed curves of bee species, reflecting within foraging bout patterns, translate into distinct abilities of bee species to move genes at the landscape level. Differences in tripping rates and in body sizes among these bee species helped explain differences among their seed curves and gene flow potential. In insect-pollinated crops, distinct bee species vary in their potential to spread genes, and future research should determine whether body size and pollinator efficiency are good general predictors of a bee species gene flow potential. The rapid development of novel genome editing technologies and the eminent release of new GE varieties, has heightened the importance of understanding how distinct bee species differentially affect gene flow, coexistence, and the spread of GE genes to sexually compatible feral and wild populations. Knowledge of the differential gene flow risk of distinct bee species can help biotechnology regulators as they refine the rules for isolation distances between fields pollinated by specific managed pollinators.

## Author Contributions

Conceived and designed the experiments: FPB JB. Set up and oversaw experiments: FPB. Analyzed the data: FPB. Funds acquisition: JB. Wrote the manuscript FPB and JB.

## Acknowledgments

We thank Molly Dieterich Mabin, Patricia Dombrowsky, Zachary Diamond and several undergraduate students for assistance with greenhouse experiments and seed tests. Luciano Palmieri assisted with the figures and Dr. Murray Clayton contributed his statistical expertise. FPF was on an appointment with the Agricultural Research Service (ARS) Research Participation Program administered by the Oak Ridge Institute for Science and Education (ORISE) through an interagency agreement between the U.S. Department of Energy (DOE) and the U.S. Department of Agriculture (USDA). ORISE is managed by ORAU under DOE [contract number DE-SC0014664]. All opinions expressed in this paper are the author’s and do not necessarily reflect the policies and views of USDA, ARS, DOE, or ORAU/ORISE.

## Data Availability Statement

Data will be made available on Dryad (www.datadryad.com) following publication.

